# The BUD13 splicing regulator: transcript structure and expression in ovules of sexual and apomictic *Paspalum notatum*

**DOI:** 10.64898/2026.06.17.732924

**Authors:** Samela Draga, Lorena A. Siena, Carolina Colono, Giovanni Gabelli, Maricel Podio, María Sol Vega, Fabio Palumbo, Juan Pablo A. Ortiz, Gianni Barcaccia, Silvina Claudia Pessino

## Abstract

**Background and Aims:** *Paspalum notatum* reproduces through either sexuality or apomixis, two pathways that may coexist within the same individual and are regulated by interconnected molecular networks responsive to environmental cues. Here, we characterized the transcript structure and expression of *BUD SITE SELECTION PROTEIN 13 (BUD13)*, a component of the RES spliceosomal complex previously reported as differentially expressed in florets of sexual and apomictic plants, as a first step toward testing its involvement in the molecular regulation of the apomixis–sexuality switch.

**Methods:** Previously generated floral and leaf transcriptomes from sexual and apomictic *Paspalum notatum* plants, including Oxford Nanopore long-read data, were mined to characterize *BUD13* transcript structure and expression. Phylogenetic analyses and in silico mapping were conducted to infer evolutionary relationships and determine the origin of the transcripts. Differential expression was validated by RT-qPCR, while *in situ* hybridization was used to reveal cell-specific ovule expression patterns.

**Key results:** *BUD13* is expressed in *Paspalum notatum* florets as a truncated isoform (*SHORT*) encoding a small protein lacking part of the herpes simplex virus regulatory protein (ICP4) domain. Two *SHORT* transcripts, *SHORT1* and *SHORT2*, with different 5′ untranslated region (UTR) regions, were identified in flowers. *SHORT1* was consistently upregulated in apomictic ovules from premeiosis to anthesis. Both transcripts originated from a single genomic locus located in the subtelomeric region of the short arm of chromosome 6. *SHORT* isoforms with variable structures were detected in other monocots. *In situ* hybridization showed that, whereas *BUD13* was expressed throughout sexual ovules, expression was absent from the female germline of apomictic ovules. A consistent expression was observed in somatic proembryos of aposporous embryo sacs.

**Conclusions:** Our findings reveal structural, spatial and temporal divergence in *BUD13* expression between sexual and apomictic reproductive programs, providing new insights into the molecular regulation of asexual seed formation.

## INTRODUCTION

Apomixis, one of the most intriguing yet underexplored phenomena in plants, is an asexual mode of reproduction that produces clonal offspring enclosed within true seeds (Hand and Koltunow, 2014; León-Martínez and Vielle-Calzada, 2019; Barcaccia *et al*., 2020). The gametophytic mechanisms underlying apomixis involve subtle deviations from the sexual reproductive pathway. These alterations occur within the ovule and lead to the formation of unreduced female gametophytes that retain the genetic constitution and ploidy level of the mother plant somatic cells (Hand and Koltunow, 2014). These atypical megagametophytes arise through the development of unreduced megaspores formed in the absence of meiosis, either directly from the megaspore mother cell (MMC), in diplosporous apomixis, or from adjacent nucellar cells, in aposporous apomixis (Hand and Koltunow, 2014). Embryo development then proceeds via parthenogenesis, without fertilization, whereas endosperm formation may occur either autonomously or following fertilization (i.e., pseudogamy), depending on the species (Hand and Koltunow, 2014). As a result, apomictic offspring inherit the full genetic complement of the mother plant, producing genetically stable clones through seeds. This trait is of great value for plant breeding and seed production because it enables the fixation of elite genotypes across generations (Barcaccia and Albertini, 2013).

Despite differences between sexual reproduction and apomixis, the two mechanisms are closely related and not mutually exclusive, as they can occur simultaneously within the same individual, either in different ovules or even within a single ovule. This coexistence is evident in facultative apomictic species, which produce heterogeneous progenies consisting of both maternal and biparental offspring (Noyes and Givens, 2013; Rodrigo *et al*., 2017; Karunarathne *et al*., 2020; Ulum *et al*., 2020; Reutemann *et al*., 2022). A major challenge in plant reproductive research is to unravel the complex molecular mechanisms linking apomixis and sexuality. This effort is supported by the increasing availability of whole-genome and tissue-specific transcriptome assemblies generated through next-generation sequencing technologies, as well as by the development of advanced genome-editing tools.

Comparative transcriptomic analyses of pistils or florets from sexual and apomictic plants have proven effective for identifying candidate genes associated with apomixis. Such studies have been conducted in a wide range of species, including *Pennisetum ciliare* (Vielle-Calzada *et al*., 1996; Singh *et al*., 2007), *Brachiaria brizantha* (Leblanc *et al*., 1997; Rodrigues *et al*., 2003; Koehler *et al*., 2020), *Panicum maximum* (Chen *et al*., 1999; Yamada-Akiyama *et al*., 2009; Radhakrishna *et al*., 2018), *Poa pratensis* (Albertini *et al*., 2004), *Eragrostis curvula* (Cervigni *et al*., 2008; Carballo *et al*., 2021; Pasten *et al*., 2022), *Paspalum notatum* (Pessino *et al*., 2001; Laspina *et al*., 2008; Ortiz *et al*., 2017, 2019; de Oliveira *et al*., 2020; Podio *et al*., 2021), *Paspalum simplex* (Polegri *et al*., 2010), *Hypericum perforatum* (Galla *et al*., 2013, 2015, 2017, 2019), *Ranunculus auricomus* (Pellino *et al*., 2013, 2020), *Hieracium* spp. (Guerin *et al*., 2000; Okada *et al*., 2013; Bräuning *et al*., 2018), and *Boechera* spp. (Sharbel *et al*., 2009, 2010; Schmidt *et al*., 2014), among others. Notably, some differentially expressed candidate genes identified in these studies map to genomic regions known to control apomixis (Schmidt *et al*., 2020).

The genus *Paspalum*, which has been the focus of apomixis research for over 50 years (Ortiz *et al*., 2013, 2020), has provided key insights into the molecular mechanisms controlling this trait. In *Paspalum*, apomixis predominantly occurs at the polyploid levels and can be either obligate or facultative (i.e., can coexist with some degree of sexuality). Mapping analyses showed that in species such as *P. notatum* and *P. simplex* the switch from sexuality to apomixis is controlled by a single genomic locus named the Apomixis Controlling Locus (ACL) (Stein *et al*., 2007; Calderini *et al*., 2011), which is transmitted with a distorted ratio. This genomic region appears to control both apospory and parthenogenesis, as well as pseudogamy. Recently, an annotated reference genome was released for *P. notatum*, facilitating the analysis of candidate genes involved in sexual and apomictic reproduction (Vega *et al*., 2024).

Early comparative transcriptomic analyses by Pessino *et al*. (2001) and Laspina *et al*. (2008) identified transcripts associated with either sexual or asexual development. A significant breakthrough came with the release of Roche 454 reference transcriptomes (Ortiz *et al*., 2017) and, later, quantitative Illumina TruSeq (MidSeq/Hiseq) transcriptomes (Ortiz *et al*., 2019; Podio *et al*., 2021). These studies revealed hundreds of differentially expressed genes between sexual and apomictic plants, several of which were subsequently functionally characterized in *P. notatum* and/or *Arabidopsis thaliana*. One such gene encodes MAP3K QUIGON JINN (QGJ), whose expression in the nucellus has been linked to the formation of aposporous embryo sacs (AES) (Mancini *et al*., 2018). In sexual ovules, *QGJ* expression is restricted to the megaspore mother cell (MMC) and later to the degenerating megaspores through the action of *TRIMETHYL GUANOSINE SYNTHASE 1* (*TGS1*) (Siena *et al*., 2014; Colono *et al*., 2019, 2022). TGS1 is specifically expressed in the proximal nucellus/chalaza/funiculus region, and its downregulation leads to ectopic *QGJ* expression in the nucellus (Colono *et al*., 2022), resulting in apospory-like phenotypes in both *P. notatum* and *A. thaliana* (Colono *et al*., 2019; Siena *et al*., 2023). Moreover, expression of *AUXIN RESPONSE FACTOR 10* (*ARF10)* in a specific cell layer surrounding the MMC is essential for proper differentiation and regular gametogenesis in *A. thaliana* (Pessino *et al*., 2024). In this model species, ectopic overexpression of *ARF10* in the nucellus leads to the formation of supernumerary reduced and unreduced functional megaspores, along with misoriented additional embryo sacs, a phenotype strongly resembling apospory (Pessino *et al*., 2024). Other candidate genes, such as the auxin-response repressor *IAA30*, are currently under investigation (Siena *et al*., 2022).

In this work, we began investigating another candidate gene for apomixis, identified through previous transcriptomic analyses: the pre-mRNA splicing factor *BUD-SITE SELECTION PROTEIN 13* (*BUD13*). *BUD13*, a known interactor of TGS1 (Hausmann *et al*., 2008), exhibits differential expression in florets of sexual and apomictic plants in at least two aposporous species, *P. notatum* and *H. perforatum* (Podio *et al*., 2021; Galla *et al*., 2019). In yeast, BUD13 is part of the RETENTION AND SPLICING (RES) complex, together with snRNP-ASSOCIATED PROTEIN 17 / INCREASED SODIUM TOLERANCE PROTEIN 3 (SNU17P/IST3P) and PRE-mRNA-LEAKAGE PROTEIN 1 (PML1P) (Wysoczanski and Zweckstetter, 2016). In plants, the *BUD13* ortholog has so far been functionally characterized only in *Arabidopsis*, where *bud13* mutants exhibit defects in early embryo development (Xiong *et al*., 2019, 2022).

The aim of this study was to characterize the *BUD13* transcript isoforms expressed in florets of sexual and apomictic *P. notatum* genotypes, analyze their phylogenetic distribution across the plant kingdom, and investigate their spatial expression patterns in ovules of both reproductive types.

## MATERIALS AND METHODS

### Plant material

The study used tetraploid (2n = 4x = 40) *P. notatum* plants, namely Q4117, a naturally occurring apomictic accession from southern Brazil (Ortiz *et al*., 1997), and Q4188, an experimentally generated fully sexual tetraploid genotype (Quarin *et al*., 2003). These materials are part of the *Paspalum* living collection maintained at Instituto de Botánica del Nordeste (IBONE-CONICET-UNNE), Corrientes, Argentina. Vegetative replicates of each genotype were grown in pots at the Instituto de Investigaciones en Ciencias Agrarias de Rosario (IICAR, CONICET-UNR), Zavalla, Argentina.

### Bioinformatic analysis

Transcript sequences classified as *BUD13* homologs were retrieved from the *P. notatum* floral transcriptome reported by Podio *et al*. (2021). Expression data were obtained from Podio *et al*. (2021). The transcripts were aligned to the *BUD13* gene annotated in the *P. notatum* reference genome (Vega *et al*., 2024) using Geneious Prime 2026.1.1 software (http://www.geneious.com). Nucleotide sequences were translated using the ExPASy Translate Tool (https://web.expasy.org/translate/), and the resulting protein sequences were aligned with Clustal Omega (https://www.ebi.ac.uk/jdispatcher/msa/clustalo). Protein domain architecture was analyzed using the NCBI Batch CD-Search tool. Custom R scripts (R Core Team, 2024) were used to classify each protein as LONG or SHORT according to its total length, using an arbitrary threshold of 480 amino acids.

### Phylogenetic analysis

A Neighbor-Joining phylogenetic tree (Saitou and Nei, 1987) was constructed with Mega12 (Sudhir *et al*., 2024). The optimal tree was constructed, showing the percentage of replicate trees, in which the associated taxa clustered together in the bootstrap test (1,000 replicates) (Felsenstein, 1985). The tree is drawn to scale, with branch lengths in the same units as those of the evolutionary distances used to infer the phylogenetic tree. The evolutionary distances were computed using the Poisson correction method (Zuckerkandl and Pauling, 1965) and are in the units of the number of amino acid substitutions per site.

### Genome mapping

Minimap2 v2.26 (Li, 2018, 2021) was used to map all the *BUD13* homologous transcripts to *P. notatum* reference genome (Vega *et al*., 2024) (NCBI accession: GCA_036689595.1), using the default parameters (-x splice).

### RT-qPCR primer design and amplification trials

For RT-qPCR assays, we manually designed the following primer pair targeting sequences exclusively present in the 5′UTR extension of the *SHORT1* isoform: BUD13A_F (forward), CCACAGAGGCGTGTGAGACAC (Tm: 61 °C), and BUD13A_R1 (reverse), CGCCTCTGTGGCAGCGAGG (Tm: 64 °C). The expected amplicon size was 113 bp. Total RNA was extracted from spikelets using the SV Total RNA Isolation Kit (Promega, Madison, WI, USA). cDNA synthesis was performed using SuperScript II (Invitrogen, Carlsbad, CA, USA) according to the manufacturer’s instructions.

Quantitative PCR reactions (final volume: 20 µL) were performed in a Rotor-Gene Q thermocycler (Qiagen, Hilden, Germany) and contained 0.5 µM gene-specific primers, 1× Real Mix qPCR (Biodynamics, Buenos Aires, Argentina), and 20 ng of cDNA. Each experiment included two biological replicates and four technical replicates per sample. The constitutively expressed gene *GLUCOSE-6-PHOSPHATE DEHYDROGENASE* (*G6PDH*) was used as the housekeeping reference gene, as described previously (Siena *et al*., 2022). Negative controls lacking template DNA were included in all experiments. Amplification was carried out under the following thermal conditions: an initial denaturation step at 94 °C for 2 min; 45 cycles of 94 °C for 15 s, 57 °C for 30 s, and 72 °C for 17 s; followed by a final extension step at 72 °C for 5 min. Gene expression levels were calculated using the ΔΔCt method (Livak and Schmittgen, 2001), assuming equal amplification efficiencies (E = 2) for both the target gene (*BUD13*) and the reference gene (*G6PDH*). Genotype Q4188 was used as the calibrator control.

### BUD13 probe amplification and labeling

A 621 bp *BUD13* fragment amplified from Q4117 cDNA was used as a probe (see previous section). This probe hybridizes to both *SHORT1* and *SHORT2* with equal specificity. To amplify the probe, the following PCR primers were designed using Primer3 v.0.4.0 (http://bioinfo.ut.ee/primer3-0.4.0/primer3/; accessed May 20, 2023): forward, AAAGGGGCAAAGGCAGTATT; reverse, ACTCAAACCATTTTTCCCCC. Total RNA was isolated using the SV Total RNA Isolation Kit (Promega, Madison, WI, USA). RNA samples were treated with RQ1 RNase-Free DNase (Promega) according to the manufacturer’s instructions to eliminate potential genomic DNA contamination. Complementary DNA (cDNA) was synthesized from 1 µg of total RNA using the ImProm-II™ Reverse Transcriptase Kit (Promega, Cat. No. A3802), following the manufacturer’s protocol. PCR amplification was performed in a T100 Thermal Cycler (Bio-Rad) in a final reaction volume of 25 µL containing 100 ng of cDNA, 1 U of Taq DNA polymerase (INBIO Highway, Tandil, Argentina), 1.5 mM MgCl₂, 0.2 µM dNTPs, and 0.2 µM of each primer. The thermal cycling program consisted of an initial denaturation step at 94 °C for 5 min, followed by 33 cycles of 94 °C for 30 s, 59 °C for 1 min, and 72 °C for 40 s. The amplified fragment was cloned into the pGEM®-T Easy vector, and the resulting recombinant plasmid was used to transform *E. coli* DH5α competent cells. Plasmids were purified using the Wizard® Plus SV Minipreps DNA Purification System (Promega), and sequence identity was confirmed by Sanger sequencing at the Aquarium Sequencing Service (Rosario, Argentina). The insert was subsequently re-amplified from the plasmid using M13 forward and reverse primers and used as a template for synthesis and DIG-labeling of sense (T7) and antisense (SP6) RNA probes using the DIG RNA Labeling Kit (SP6/T7) (Roche, Basel, Switzerland).

### *In situ* hybridization experiments

Spikelets from Q4117 and Q4188 were collected at three developmental stages—premeiosis, meiosis, and postmeiosis—according to the developmental calendar described by Laspina *et al*., (2008). Developmental stages were determined based on pollen development by crushing the anthers, staining them with 45% acetic carmine, and examining them under a light microscope. Samples were dehydrated through an ethanol/xylene series, fixed in 4% paraformaldehyde and 0.25% glutaraldehyde, embedded in paraffin, and sectioned using a Minot microtome to obtain cross or sagittal sections (10 µm thick), which were mounted on slides previously coated with poly-L-lysine (100 µg mL⁻¹). Paraffin was removed using the same ethanol/xylene series. Prehybridization was performed at 37 °C for 10 min in 0.05 M Tris–HCl buffer (pH 7.5) containing 1 µg mL⁻¹ proteinase K. Hybridization was carried out for 12 h at 37 °C in a buffer containing 10 mM Tris–HCl (pH 7.5), 300 mM NaCl, 50% deionized formamide, 1 mM EDTA (pH 8), 1× Denhardt’s solution, 10% dextran sulfate, 600 ng mL⁻¹ total RNA, and 60 ng of the corresponding probe. Signal detection was performed according to the instructions provided with the DIG detection kit (Roche, Basel, Switzerland), using anti-DIG-AP and NBT/BCIP as substrates. A Nikon Eclipse E200 microscope (Nikon, Tokyo, Japan) equipped with a digital camera was used for observation. For each experiment, three independent slides containing five to seven ovules from both apomictic and sexual genotypes were hybridized simultaneously in the same chamber. Signal development was monitored synchronously. Only hybridization patterns showing consistent signal distribution were considered. To ensure reproducibility, at least five ovules per slide (3 × 5 = 15 ovules in total) were successfully analyzed.

## RESULTS

### Selection of *BUD13* as an apomixis candidate gene

Understanding the complex molecular cascade underlying the switch from sexual to apomictic reproduction requires the identification of candidate genes suitable for functional analysis. The criteria commonly used for candidate gene selection are differential expression, association with previously characterized regulators of apomixis, and/or localization within the ACL. In this study, comparison of differentially expressed transcripts between sexual and apomictic plants in two model aposporous species—*P. notatum* (Podio *et al*., 2021) and *H. perforatum* (Galla *et al*., 2019)—identified *BUD-SITE SELECTION PROTEIN 13* (*BUD13*) as a candidate gene of particular interest. *BUD13* encodes a pre-mRNA splicing factor that is a component of the RES complex (Wysoczanski and Zweckstetter, 2016).

BUD13 attracted our attention because it has been reported to interact with TGS1 (Hausmann *et al*., 2008), a well-established apomixis-related protein (Siena *et al*., 2014, Colono *et al*., 2019; Siena *et al*., 2023). According to Hausmann *et al*. (2008), both genes participate in partially redundant pathways that support efficient pre-mRNA splicing during specialized developmental programs. The *BUD13* transcript was consistently differentially expressed between apomictic and sexual genotypes in *P. notatum* and *H. perforatum* (Podio *et al*., 2021; Galla *et al*., 2019). In *Paspalum*, it exhibited log fold changes of approximately 4 (corresponding to ∼32-fold differences in expression) across all reproductive stages analyzed, from premeiosis to anthesis (Podio *et al*., 2021).

### Structure of the *BUD13* transcript variants expressed in *Paspalum* reproductive organs

The canonical *BUD13* transcript sequence described in *A. thaliana* (AT1G31870) and other species—hereafter referred to as *LONG*—encodes a full-length protein containing a domain similar to the C-terminal region of the herpes simplex virus regulatory protein ICP4 (PHA03307, ICP4-C domain) (Bruce and Wilcox, 2002), as well as a BUD13-specific domain (Pfam09736). In the *P. notatum* genome, the *LONG* transcript isoform is encoded by a single gene (predicted locus: PnR1_U9M48_029946) (Vega *et al*., 2024), which is expressed in leaves of diploid and tetraploid sexual genotypes as well as tetraploid apomictic genotypes (Marino *et al*., 2024; transcripts DAWXEG010026322.1, DAWXED010027222.1, DAWXEG010152668.1, and DAWXEG010152667.1). Moreover, the *LONG BUD13* isoform was also detected, albeit at low levels, in *P. notatum* florets, as the complete sequence was successfully assembled from Roche 454 libraries generated to produce reference floral transcriptomes of sexual and apomictic plants (Ortiz *et al*., 2017; transcripts GFNR01015542.1 and GFMI02018658.1). Transcriptome surveys revealed additional shorter *BUD13* transcript isoforms—hereafter referred to as *SHORT*—expressed in *P. notatum* florets (Podio *et al*., 2021; transcripts GIUR01028184.1 and GIUR01125423.1). According to the transcriptome dynamics analysis reported by Podio *et al*. (2021), these *SHORT* isoforms are expressed in florets throughout reproductive development (premeiosis, meiosis, postmeiosis, and anthesis. Structurally, *SHORT* isoforms consist of truncated sequences lacking a substantial portion of the sequence encoding the ICP4 domain. Interestingly, these two *SHORT* transcripts, hereafter designated as *SHORT1* and *SHORT2,* differ in two major respects: i) the length of their 5′ UTRs, with *SHORT1* harbouring an extended 5′ UTR relative to *SHORT2*; and ii) their expression profiles in floral *P. notatum* transcriptomic data, where *SHORT1* displays an overexpression in apomictic genotypes, whereas *SHORT2* shows higher expression in sexual genotypes (see below).

*SHORT1* and *SHORT2* correspond to GenBank accessions GIUR01028184.1 and GIUR01125423.1 with transcript identifiers TRpn_125370 and TRpn_33688, respectively (Podio *et al*., 2021). A schematic representation generated by mapping the *LONG* and *SHORT* transcript isoforms onto the *P. notatum* reference genome (Vega *et al*., 2024), comparatively illustrating transcript structure, including 5′ and 3′ UTRs, exons, and introns, is shown in Fig. 1A.

**Fig. 1.**
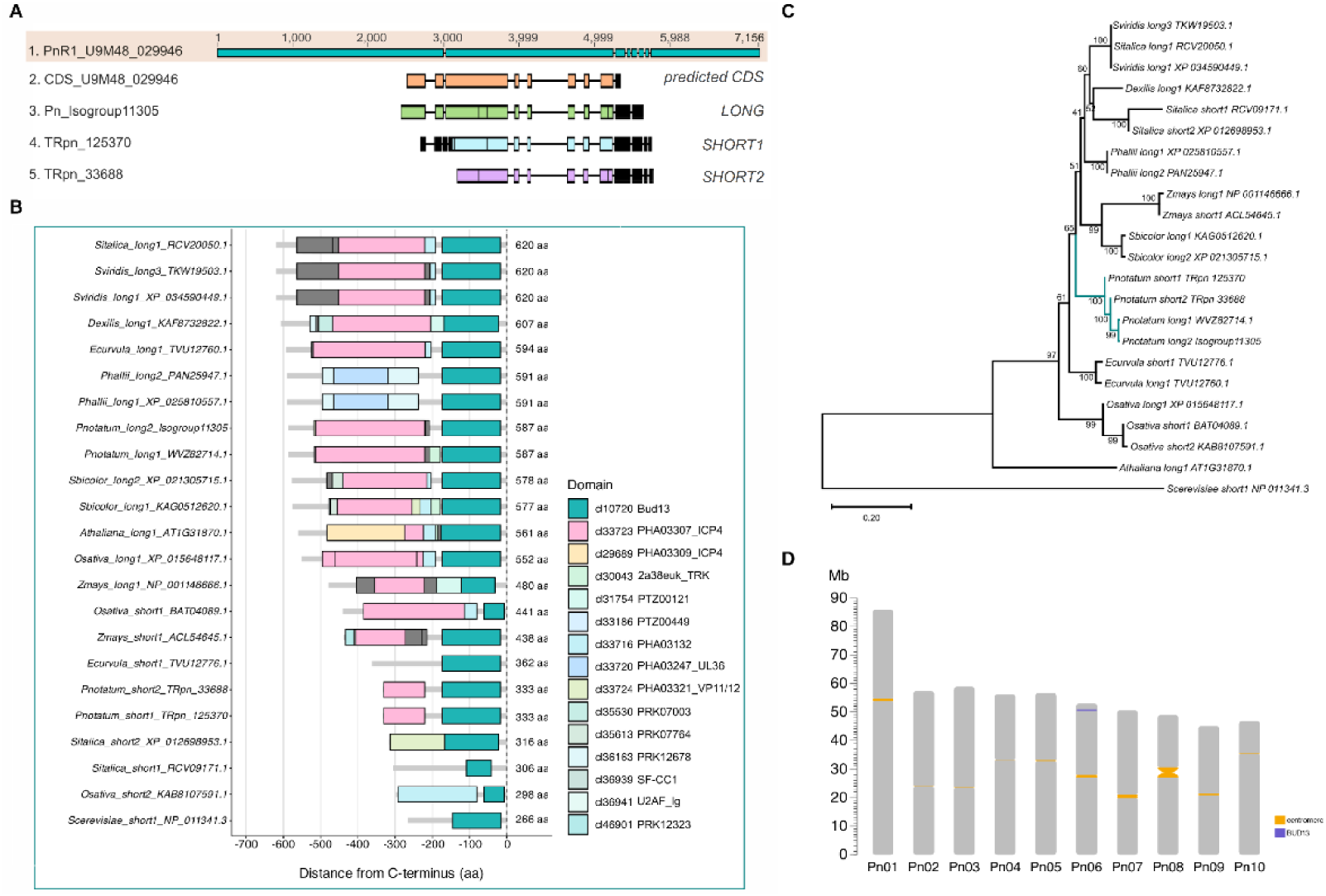
BUD13 transcript/protein structure and phylogeny. (A): Alignment of different *BUD13* transcript isoforms detected in *P. notatum* onto the R1 reference genome (PnR1_U9M48_029946) (Vega et al., 2024). The genomic DNA model (top) is shown alongside the corresponding cDNA/transcript models (bottom). CDS_U9M48_029946 corresponds to the predicted coding sequence for this genomic region (Vega et al., 2024). Colored boxes indicate exons and lines indicate introns. Black boxes represent untranslated regions (5′ or 3′ UTRs). (B): Schematic representation of the different protein domain architectures detected in BUD13 proteins from related Poaceae species and two outgroup species (*A. thaliana* and *S. cerevisiae*), ordered by protein length. Colored boxes indicate conserved domains identified in each sequence. Protein lengths (in amino acids) are shown on the right and are aligned relative to the C-terminus. (C): Neighbor-Joining phylogenetic tree constructed using the same protein sequences analyzed in panel B. The optimal tree (sum of branch lengths = 2.749) was generated, and bootstrap support values based on 1,000 replicates are indicated at each node. The analysis included 674 aligned positions after deletion of ambiguous sites by pairwise comparison. The tree was drawn to scale. As expected, the outgroup species (*A. thaliana* and *S. cerevisiae*) clustered separately from the remaining species, all of which belong to the Poaceae family. In general, LONG and SHORT isoforms from the same species clustered together, supporting the hypothesis that they represent splice variants derived from a single genomic locus. (D): In silico mapping of BUD13 homologs across the *P. notatum* reference genome (Vega et al., 2024). Bars represent chromosome sizes (Mb). The position of the *BUD13* gene is highlighted (purple line). The centromeres are indicated with yellow marks.

The existence of both *LONG* and *SHORT* transcripts was further validated through their direct identification as raw reads (e4da906b-7e29-4180-a1d0-485859108b8e and c38fedb6-e5e5-4848-a700-d242bbf39209) in an Oxford Nanopore Technology (ONT) floral transcriptome dataset generated for *P. notatum* (Vega *et al*., 2024; NCBI accession number: SRR27335061).

From the protein prediction analysis, it was observed that the *LONG* transcript isoform encodes a 587-amino-acid protein containing the complete ICP4 (PHA03307, 293 amino acids) and BUD13 (156 amino acids) domains. In contrast, both truncated *SHORT* transcript isoforms *SHORT1* (TRpn_125370) and *SHORT2* (TRpn_33688) encode a 333-amino-acid protein containing only 110 of the 293 amino acids constituting the ICP4 domain (i.e., 183 amino acids of the ICP4 domain are absent), while retaining the complete BUD13 domain (156 amino acids). Although *SHORT1* (overexpressed in apomictic genotypes) includes a 5′ UTR extension absent from *SHORT2*, both *SHORT* transcripts encode the same protein. An alignment of the full-length BUD13 protein WVZ82714.1 (as predicted from genomic locus PnR1_U9M48_029946; Vega *et al*., 2024), the Roche 454-derived isoform (isogroup11305), and the two SHORT-derived proteins (SHORT1 and SHORT2) is presented in Supplementary Fig. S1.

### BUD13 protein domain architecture and phylogenetic analysis

To investigate the presence of *LONG* and *SHORT BUD13* isoforms in plants, we performed a structural analysis of 110 BUD13 protein sequences available in public databases, originating from 40 monocot species, including *P. notatum*. Two outgroup species (*A. thaliana* and *Saccharomyces cerevisiae*) were also included in the analysis.

First, a list of all available BUD13 protein sequences from monocot species was compiled from Entrez by searching the gene name (Supplementary Data S1, page 1). Protein domain architecture was then analyzed using the NCBI Batch CD-Search tool (identified domains are listed in Supplementary Data S1, page 2). Finally, custom R scripts (R Core Team, 2024) were used to classify each protein as LONG or SHORT based on its total length (the classification is provided in Supplementary Data S1, page 2). An arbitrary threshold of 480 amino acids was applied to define proteins as LONG (>480 amino acids) or SHORT (<480 amino acids). When multiple protein sequences were present within the same species, variants were numbered consecutively (e.g., LONG1, LONG2, etc., or SHORT1, SHORT2, etc.) (Supplementary Data S1, page 2).

Among the 40 monocot species analyzed, 27 display only the LONG BUD13 protein, 9 both SHORT and LONG isoforms, and 4 only SHORT isoforms (Supplementary Data S1, page 2). The outgroup species *A. thaliana* and *S. cerevisiae* have only LONG and only SHORT isoforms, respectively. The LONG and SHORT protein sequences detected exhibit a remarkable diversity in domain composition, suggesting functional diversification across species (Supplementary Data S1, page 2). Out of the 110 protein sequences analyzed, we selected 21 originating from the Gramineae, representing different species and isoforms, plus the sequences from the two outgroup species, to construct a diagram illustrating the diversity of domain combinations (Fig. 1B). Most SHORT sequences have truncated or absent ICP4 domains; however, some retain a complete ICP4 domain and a truncated BUD13 domain, such as *O. sativa*_short1_BAT04089.1 (Fig. 1B).

To explore the evolutionary relationships among species, a phylogenetic analysis was conducted using the same 110 BUD13 protein sequences from monocots previously employed for structural analysis (Supplementary Fig. S2).

As expected, outgroup sequences clustered separately from the remaining sequences. *Carex littledalei* (Cyperaceae) also clustered separately from the rest of the monocots. The remaining sequences grouped into two main clades: (1) a major clade containing all sequences originating from the Poaceae, including *P. notatum*; and (2) a minor clade comprising sequences originating from Bromeliaceae, Araceae, Orchidaceae, Asparagaceae, Arecaceae, Dioscoreaceae, Musaceae, Zingiberaceae, and Zosteraceae.

For species with reported LONG and SHORT protein variants in databases (*Oryza sativa*, *P. notatum*, *Hordeum vulgare*, *Zea mays*, *E. curvula*, *Triticum turgidum*, *Setaria italica*, *Asparagus officinalis*, *Zingiber officinale*), both protein variants clustered together within the same phylogenetic group suggesting a common origin.

We also constructed an analogous phylogenetic tree using the 21 *Poaceae* protein sequences employed for protein domain characterization (Fig. 1C). As expected, the *S. cerevisiae* sequence (a SHORT isoform) was positioned as an outgroup, while the *A. thaliana* sequence (a LONG isoform) was also located outside the Poaceae cluster. The remaining sequences grouped into clusters according to species identity; that is, LONG and SHORT sequences from the same species clustered together. The only exception to this pattern was observed in *S. italica* and *Setaria viridis*, where variation between SHORT and LONG appeared to outweigh species-specific variation in defining cluster formation. It should be noted that *S. viridis* is the wild progenitor of *S. italica*; therefore, the genetic distance between these two species is expected to be minimal (Fig. 1C).

In general, these results indicate that, within a given species, LONG and SHORT protein sequences are highly similar and likely originate either from a single gene (i.e., as alternative splice variants) or from recently duplicated genes that have not yet diverged through mutation. Supporting this hypothesis, when the *LONG* and *SHORT* (*SHORT1* and *SHORT2*) *BUD13* transcripts were mapped onto the diploid *P. notatum* reference genome (Fig. 1D), all aligned to a single genomic region corresponding to the *BUD13* locus, located in the subtelomeric region of the short arm of chromosome 6. As this is the only genomic region homologous to *BUD13* in the *Paspalum* genome, the *LONG* and *SHORT* transcripts most likely represent splice variants or transcripts originating from alternative transcription start sites (TSSs) of the *BUD13* gene.

### Sequencing-based quantification and RT-qPCR validation of *SHORT1* upregulation in apomictic ovules

Differential expression analyses between sexual and apomictic floral transcriptomes reported by Podio *et al*. (2021) revealed that *SHORT1* is upregulated in apomictic plants from premeiosis to anthesis, reaching peak expression at the latter developmental stage, whereas *SHORT2* expression remains consistently low through all reproductive stages (Fig. 2). The opposite pattern was observed in sexual plants: *SHORT2* is overexpressed throughout development, reaching its highest expression level at anthesis, while *SHORT1* expression remains low (Fig. 2).

**Fig. 2.**
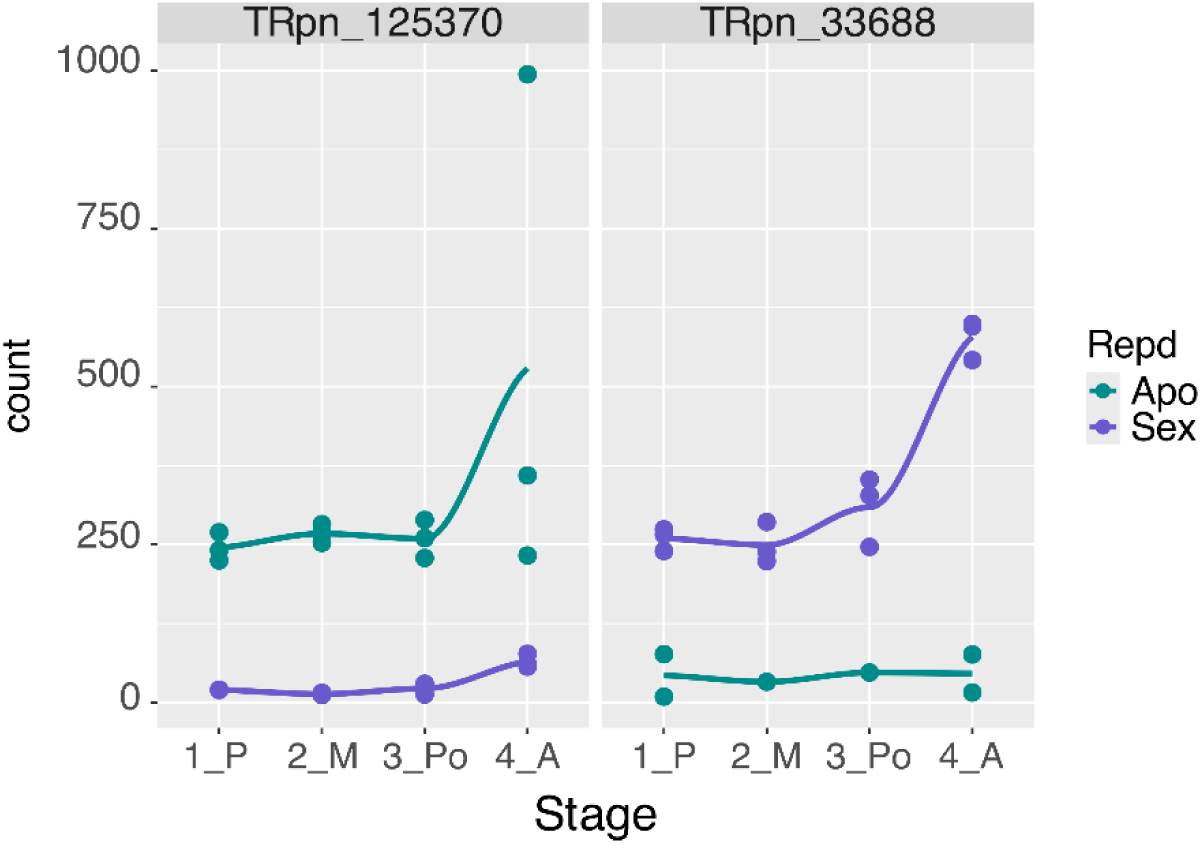
Progression of *SHORT1* and *SHORT2* expression during reproductive development in sexual and apomictic ovules. Left panel: expression of TRpn_125370 (*SHORT1*) in apomictic (green lines) and sexual (purple lines) genotypes at premeiosis (1_P), meiosis (2_M), postmeiosis (3_Po), and anthesis (4_A). Right panel: expression of TRpn_33688 (*SHORT2*) in apomictic (green lines) and sexual (purple lines) genotypes at premeiosis (1_P), meiosis (2_M), postmeiosis (3_Po), and anthesis (4_A). Expression of both floral *SHORT* transcript variants increased at anthesis. The replicate dispersion observed for *SHORT1* expression at anthesis in apomictic ovules may be associated with the abrupt increase in transcript accumulation detected at this developmental stage.

Taking advantage of the 5′ UTR extension present in the *SHORT1* sequence, which is absent in *SHORT2*, we designed specific primers to exclusively amplify this isoform in RT-qPCR assays (Fig. 3A, B), in order to validate the upregulation detected for this isoform in florets of apomictic plants. We found that, at premeiosis/meiosis, *SHORT1* was overexpressed by a factor of 18.8 ± 6.747 (*t-*test, *p* = 0.038) in florets of apomictic genotypes compared to sexual ones (Fig. 3C). These results are consistent with the sequencing analysis reported by Podio *et al*. (2021).

**Fig. 3.**
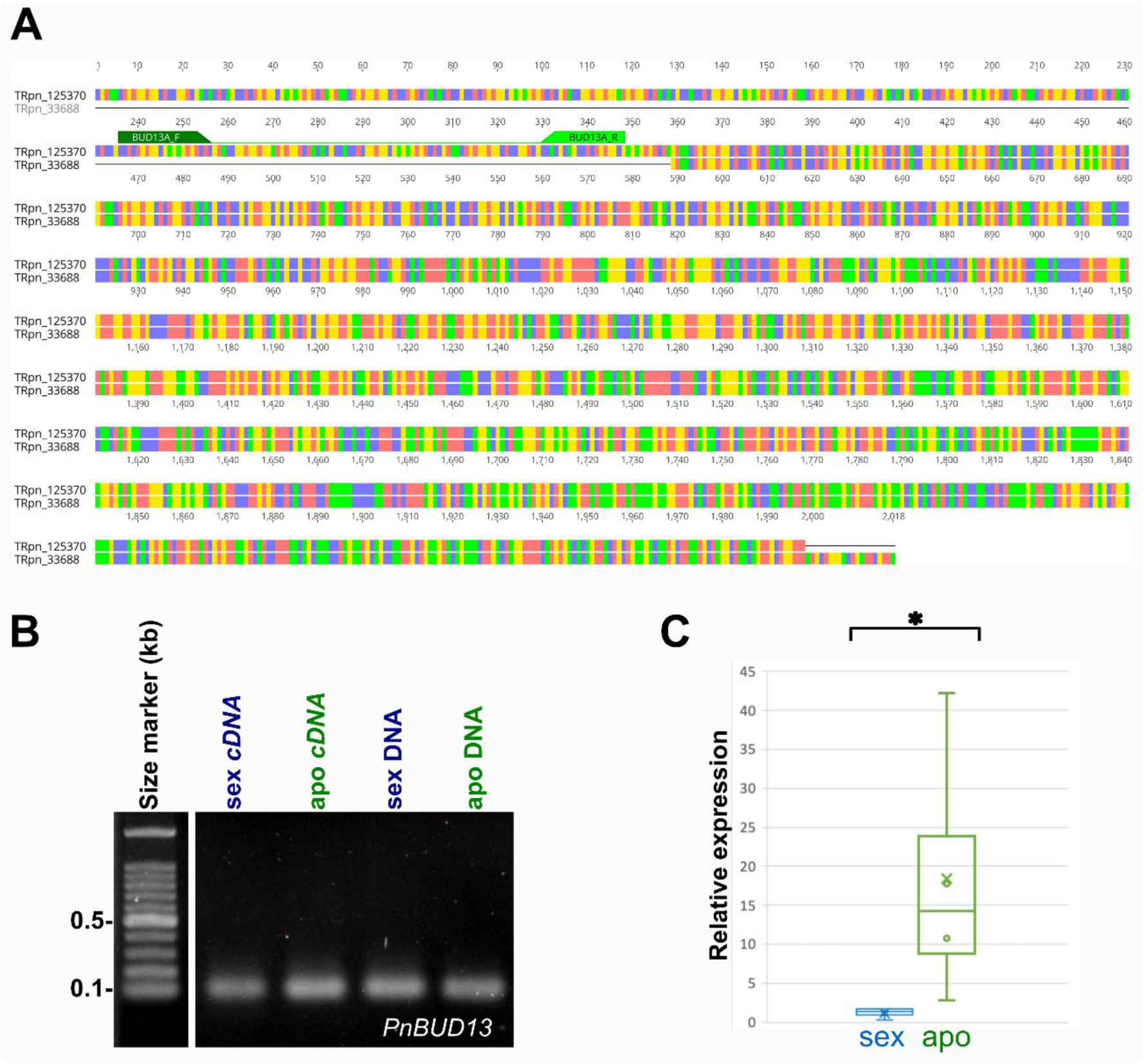
Expression of the BUD13 isoform SHORT1 at premeiosis in florets of sexual and apomictic plants. (A): Comparison of the nucleotide sequences of *SHORT1* and *SHORT2* revealed that *SHORT1* contains an extended 5′ untranslated region (5′ UTR). Specific qPCR primers (BUD13A_F and BUD13A_R) were designed to target this region. (B): Non-quantitative PCR amplification of floral cDNA at premeiosis using BUD13A_F and BUD13A_R yielded a single amplicon of the expected size (113 bp). The sequence was detected in both sexual and apomictic plants. (C): qPCR analysis at premeiosis revealed statistically significant differential expression of *SHORT1* between apomictic and sexual samples. Boxplots show the interquartile range (25th–75th percentiles), the median (line within the box), and the mean (×). Upper and lower whiskers indicate the 95th and 5th percentiles, respectively, and outliers are represented by circles. * indicates significant differences at p < 0.05. Nucleotide color code: red, adenine; blue, cytosine; yellow, guanine; green, thymine.

### BUD13 shows differential *in situ* distribution patterns in sexual and asexual *P. notatum* plants

To gain further insight into BUD13 expression, we analyzed its spatial distribution in reproductive organs by performing *in situ* hybridization assays on ovaries containing ovules at the premeiosis/meiosis and anthesis stages in both apomictic and sexual genotypes. We could not use a probe specific for *SHORT1* because the fragment specifically amplified from this isoform in the RT-qPCR experiments contains internal repetitive sequences shared by both *SHORT* isoforms (*SHORT1* and *SHORT2*). Therefore, we selected a probe equally specific for both isoforms. Accordingly, we inferred that the hybridization pattern detected in ovules of apomictic and sexual plants reflects the expression of the TRpn_125370 and TRpn_33688 sequences, respectively, as documented in Illumina sequencing experiments (Podio *et al*., 2021) and in the RT-qPCR validation assays.

Hybridization with the antisense probe (SP6, detecting the coding strand) performed on ovules prior to female meiosis (MMC stage) revealed signals restricted to the nucellus and integuments in both sexual (Fig. 4A, C) and apomictic genotypes (Fig. 4D, F), contrasting with the surrounding somatic tissues. However, the megaspore mother cell (MMC) exhibited a consistent (n = 8) differential pattern between reproductively contrasting genotypes. In sexual ovules, the MMC displayed a staining intensity comparable to that of adjacent nucellar cells (Fig. 4C), whereas in apomictic ovules, no signal was detected in either the MMC or the aposporous initial cells (Fig. 4F). In anthers, microspore mother cells showed detectable signals in both genotypes (Fig. 4B, E), although they were stronger and more uniform in sexual plants. A similar hybridization in observed in the tapetum cells. Control hybridizations using a sense probe (T7, detecting the antisense transcript strand) showed no signal in the MMCs, initial cells, nucellar cells, or integuments (Fig. 4G, I, J, L), as well as in microspores and tapetal tissue (Fig. 4H, K) of both genotypes.

**Fig. 4.**
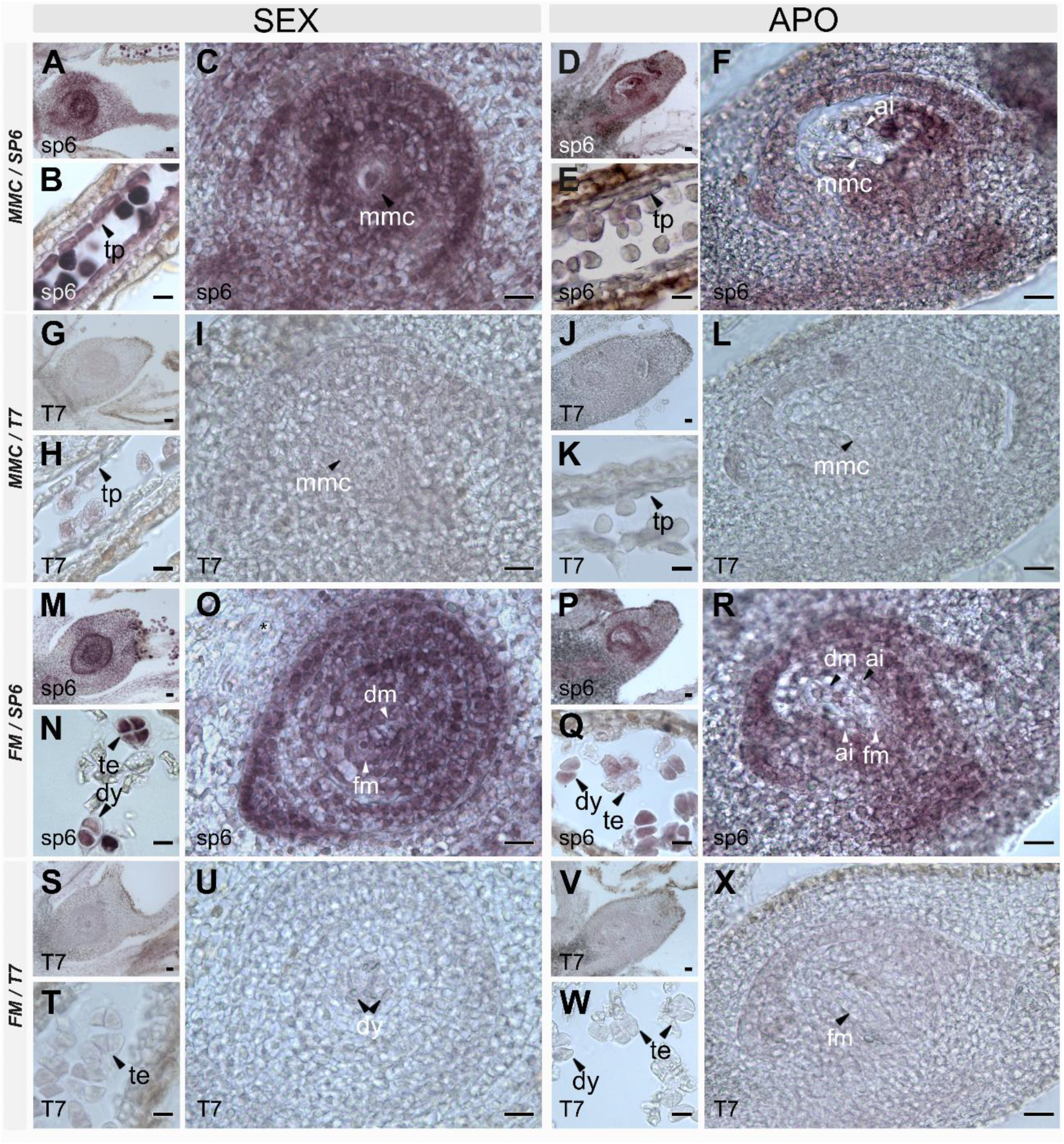
RNA *in situ* hybridization of BUD13 transcripts in reproductive organs of sexual (Q4188) and apomictic (Q4117) *Paspalum notatum* at premeiosis/meiosis. (A–F): Hybridization with the antisense probe (SP6) in cross or sagittal sections of ovules at the megaspore mother cell (MMC) stage and anthers showing the tapetum and pollen mother cells in sexual (A–C) and apomictic (D–F) plants. (G–L): Hybridization with the sense probe (T7) in sagittal sections of ovules at the MMC stage and anthers with tapetum and microspore mother cells in sexual (G–I) and apomictic (J–L) individuals. (M–R): Hybridization with the antisense probe (SP6) in cross or sagittal sections of ovules at the functional megaspore (FM) stage and anthers containing microspore tetrads in sexual (M–O) and apomictic (P–R) plants. (S–X): Hybridization with the sense probe (T7) in cross or sagittal sections of ovules at the FM stage and anthers with microspore tetrads in sexual (S–U) and apomictic (V–X) individuals. Abbreviations: ai: apospory initials; apo, ovules from Q4117; dy, dyad cells; dm, degenerating megaspores; fm, functional megaspore; mmc, megaspore mother cell; tp, tapetum; te, tetrads; sex, ovules from Q4188. Bars: 10 μm.

Analysis of hybridization patterns using the antisense probe (SP6) at the functional megaspore (FM) stage also revealed signals in the nucellar cells and integuments of both sexual and apomictic genotypes (Fig. 4M, O, P, R). Consistent with the previous stage, the FM and degenerating megaspores of sexual ovules displayed hybridization levels comparable to those of the surrounding somatic cells (Fig. 4O), whereas the megaspores and adjacent aposporous initial cells in apomictic ovules showed no detectable signal (Fig. 4R). In anthers, microspores of both genotypes exhibited hybridization signals (Fig. 4N, Q) but, as previously observed, the signal was stronger and more uniform in the sexual genotype. Hybridization with the sense probe (T7) once again revealed no antisense transcript signal in either genotype (Fig. 4S–X).

Additional images provided in Supplementary Fig. S3 illustrate the *BUD13* expression in sexual genotypes compared to apomictic ones at the functional megaspore stage.

At anthesis, hybridization with the antisense probe (SP6) again revealed a contrasting expression pattern between sexual and apomictic genotypes. In sexual ovules, strong hybridization signals were detected in nucellar tissues, integuments and reproductive cells (Fig. 5A, B), whereas in apomictic ovules, only weak signals were observed in nucellar cells, egg cells, synergid cells, and polar nuclei (Fig. 5D, E, G, H, J, K). In adjacent sections of the same ovule, no signal was detected in antipodal cells (Fig. 5E, G, H). Notably, *BUD13* expression was detected only in a subset of egg cells from different aposporous embryo sacs within the same ovary (compare Fig. 5H, K), suggesting a stage-specific activation of this gene in that cell type. The sense probe (T7) confirmed the absence of antisense transcript signal in both sexual (Fig. 5C) and apomictic genotypes (Fig. 5F, I, L) at this stage. Additional images illustrating the hybridization pattern at anthesis are provided in Supplementary Fig. S4 and S5. Further images demonstrating the absence of antisense transcript signal using the sense probe (T7) are shown in Supplementary Fig. S6.

**Fig. 5.**
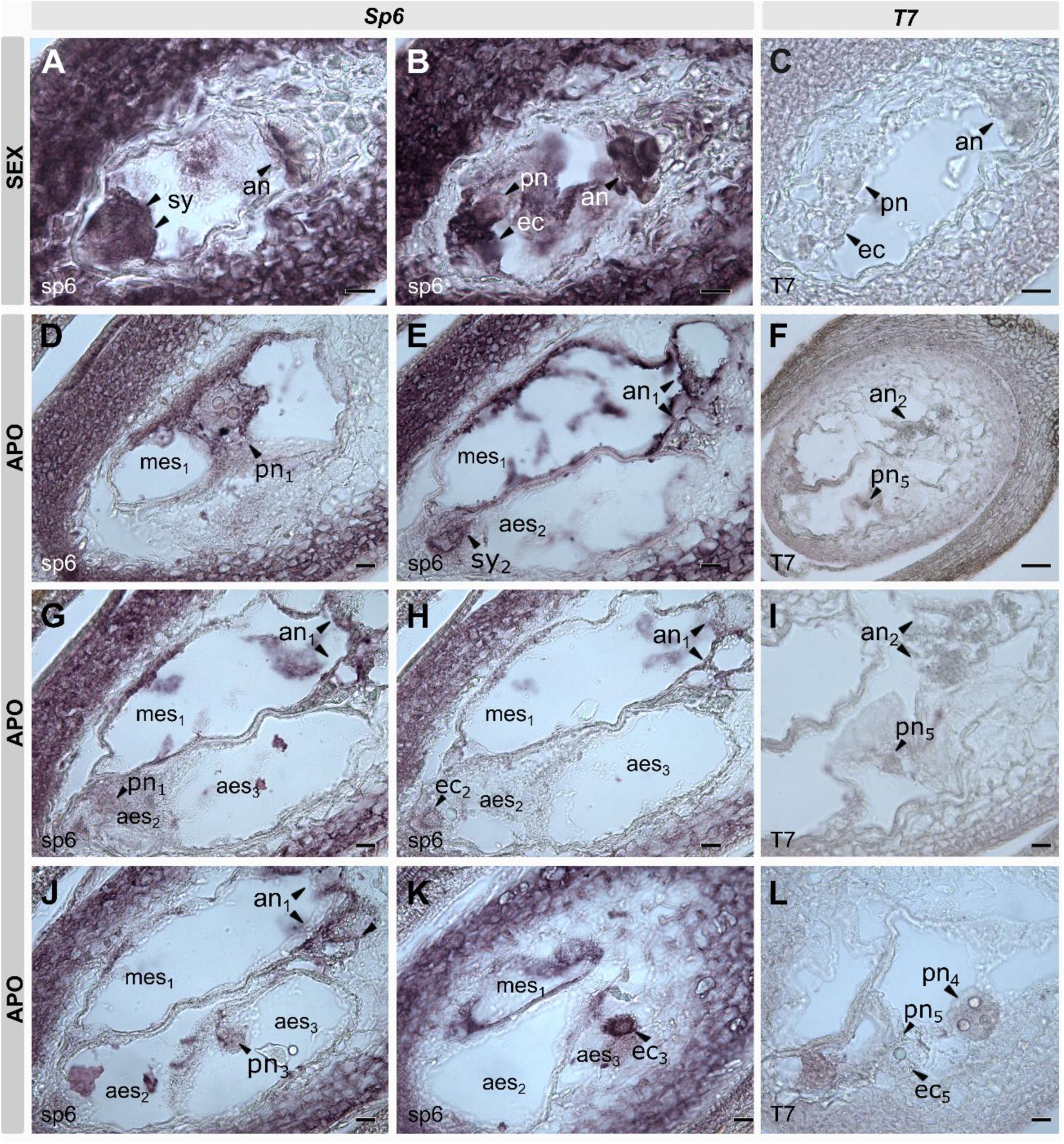
RNA *in situ* hybridization of BUD13 transcripts in ovaries from sexual (Q4188) and apomictic (Q4117) *Paspalum notatum* at anthesis. (A–C): Sexual genotype. Hybridization with the *BUD13* antisense probe (SP6) (A, B), and with the sense probe (T7) (C). (D–L): Apomictic genotype. Hybridization with the *BUD13* antisense probe (SP6) (D, E, G, H, J, K; adjacent sections of the same ovule) and with the sense probe (T7) (F, I, L; adjacent sections of the same ovule). Abbreviations: an, antipodal cells; ec, egg cell; aes, aposporic embryo sac; mes, meiotic embryo sac; pn, polar nuclei; sy, synergid cells. The subscript number indicates the embryo sac in which the structure is located. Bars: 10 μm.

Notably, in apomictic ovules at anthesis, parthenogenetic proembryos displayed hybridization signals similar to those observed in integumental cells (Fig. 6A, B). These proembryos develop from aposporous embryo sacs, which lack antipodal cells. Hybridization with the sense probe did not reveal any signal (Fig. 6C, D). Finally, no NBT/BCIP staining was detected in any of the negative controls (data not shown).

**Fig. 6.**
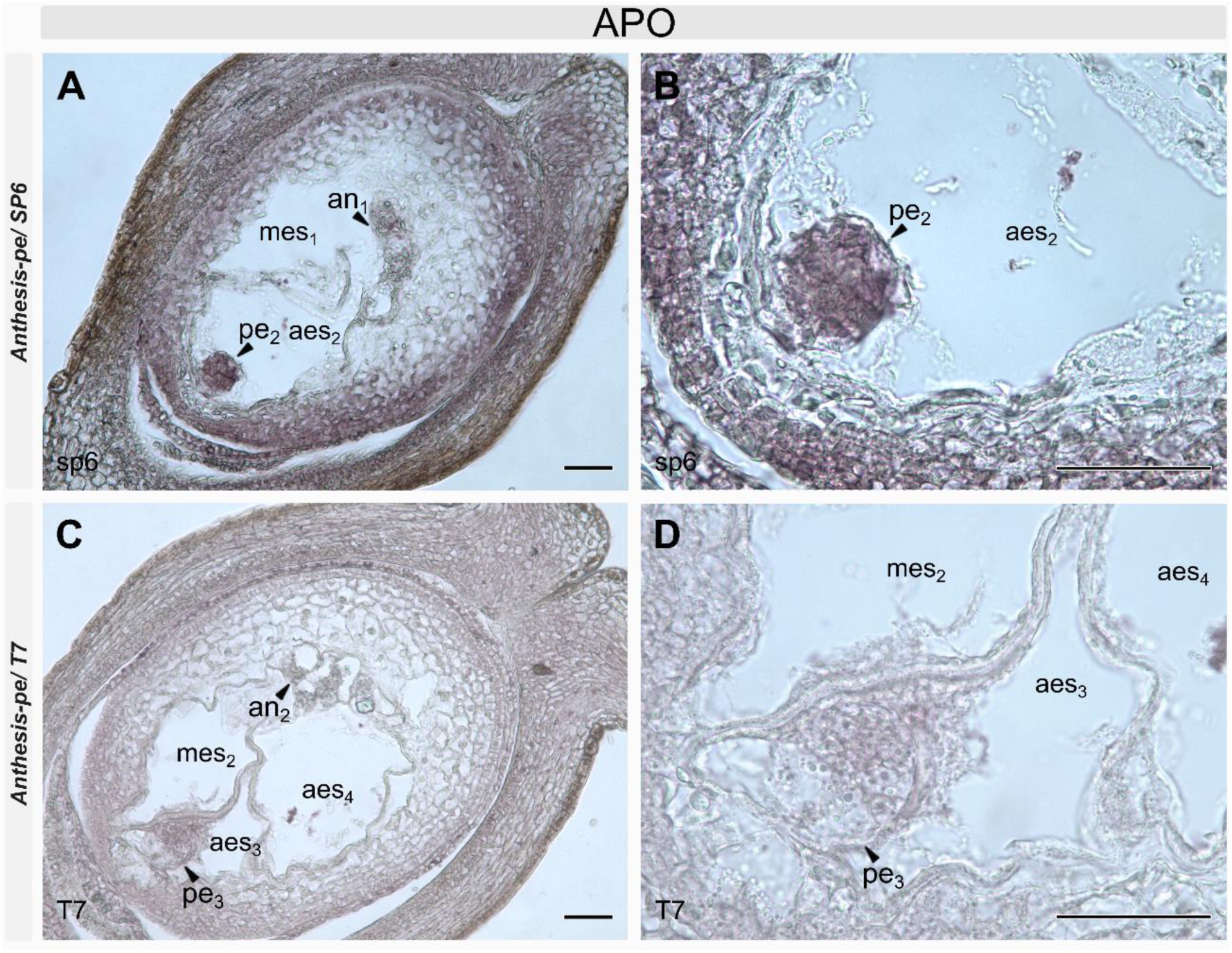
RNA *in situ* hybridization of BUD13 transcripts in proembryos from apomictic ovaries (Q4117) of *Paspalum notatum*. (A, B): Hybridization with the antisense probe (SP6). (C, D): hybridization with the sense probe (T7), shown at different focal planes. Note the presence of multiple embryo sacs, indicated by subscript numbers, and the parthenogenetic embryos (pe₂ in A, B; pe₃ in C, D) within some of the aposporous embryo sacs, but not in the meiotic sacs. Abbreviations: an, antipodal cells; aes, aposporic embryo sac; mes, meiotic embryo sac; pe, proembryo. The subscript number indicates the embryo sac in which the structure is located. Bars: 10 μm.

## DISCUSSION

In recent years, apomictic development has been functionally linked to the RNA splicing machinery through the identification of TRIMETHYLGUANOSINE SYNTHASE 1 (TGS1) as a major inhibitor of aposporous development (Siena *et al*., 2014, 2023; Colono *et al*., 2019). TGS1 is an RNA-binding protein containing a methyltransferase domain that performs a dual function as both a transcriptional regulator and a controller of RNA processing. Its methyltransferase activity converts canonical m7G caps into the 2,2,7-trimethylguanosine (TMG) caps characteristic of spliceosomal small nuclear RNAs. Hypermethylated 2,2,7-trimethylguanosine (TMG) cap structures are characteristic of small nuclear and nucleolar RNAs involved in pre-mRNA splicing (U1, U2, U4, and U5), pre-rRNA processing (U3 and U8), and telomere addition (telomerase RNA) (Seto *et al*., 1999; Liou and Blumenthal, 1990). In *P. notatum*, *TGS1* is expressed at lower levels in the ovules of apomictic plants than in those of sexual plants (Siena *et al*., 2014). Sexual plants with reduced *TGS1* expression generated through antisense technology exhibit misoriented supernumerary embryo sacs resembling those observed in apomictic individuals (Colono *et al*., 2019). However, neither proembryos nor apomixis-derived seeds were detected in these plants, suggesting that *TGS1* is required to repress the formation of aposporous embryo sacs but is not sufficient to trigger parthenogenesis (Colono *et al*., 2019). Moreover, A. thaliana tgs1-defective mutants exhibited altered specification of the precursor cells that give rise to the female gametophyte and the sporophyte, resulting in the formation of a functional aposporous-like cell lineage and further supporting the involvement of this candidate gene in apospory (Siena *et al*., 2023).

The TGS1 interaction network was characterized in yeast and consists of proteins involved in small nuclear ribonucleoprotein function and spliceosome assembly, including MUD2, NAM8, BRR1, LEA1, IST3, ISY1, CWC21, and BUD site selection protein 13 (Bud13p) (Hausmann *et al*., 2008). Particularly, Bud13p is part of the RES complex, a 71 kDa heterotrimer composed of the proteins Pre-mRNA leakage protein 1 (Pml1p), U2 snRNP component Snu17 (Snu17p) and Bud13p (Wysoczanski and Zweckstetter, 2016). Yeast Bud13p interacts directly with Snu17p (Wysoczanski and Zweckstetter, 2016), but not with Pml1p (Wysoczanski and Zweckstetter, 2016; Trowitzsch *et al*., 2008; Brooks *et al*., 2009; Wysoczanski *et al*., 2014, 2015). The binding of Bud13p to Snu17p promotes the interaction between Snu17p and Pml1p (Wysoczanski *et al*., 2014). In the absence of Bud13p, the splicing efficiency of reporter constructs is substantially decreased (Gottschalk *et al*., 2001; Dziembowski *et al*., 2004). So far, the splicing of four pre-mRNAs—ACT1, TAN1, MER1, and MATa1—has been shown to be controlled by Bud13p (Wysoczanski and Zweckstetter, 2016; Clark *et al*., 2002; Khanna *et al*., 2009; Zhou *et al*., 2013; Xu *et al*., 2018). In *A. thaliana*, AtBUD13 knockout mutations lead to defective splicing of 52 genes associated with early embryo development, resulting in embryo lethality (Xiong *et al*., 2019). Moreover, Xiong et al. (2022) determined that *A. thaliana* BUD13 (AtBUD13), GROWTH, DEVELOPMENT AND SPLICING 1 (GDS1), and DAWDLE (DDL) are the respective counterparts of the yeast RES complex subunits Bud13p, Snu17p, and Pml1p.

Since both the well-characterized apospory regulator *TGS1* and *BUD13* participate in shared interaction networks associated with RNA splicing, we decided to explore a possible role for the latter in apomictic development. Indeed, *BUD13* has been reported to be differentially expressed in the floral transcriptomes of sexual and apomictic plants of *P. notatum* (Podio *et al*., 2021) and *H. perforatum* (Galla *et al*., 2019). We retrieved the transcript sequences expressed in *P. notatum* florets and identified a short *BUD13* transcript isoform encoding a small protein (BUD13_SHORT) that contains the BUD13 domain but lacks part of the ICP4-C domain, in contrast to the canonical isoform, which contains both complete domains. Moreover, transcripts encoding this *SHORT* isoform occur in two variants, *SHORT1* and *SHORT2*, with and without a terminal 5′ UTR extension, respectively. These variants are derived from the same genomic locus and are overexpressed in apomictic and sexual plants, respectively. Since the 5′ UTRs of messenger RNAs (mRNAs) regulate translation, participate in ribosome recruitment and progression, influence translation efficiency, mediate responses to environmental changes, and affect transcript stability, the presence of two *SHORT BUD13* transcript variants differing in their 5′ UTR structure in apomictic and sexual genotypes may modulate one or more of these processes, potentially impacting development.

Phylogenetic analysis based on monocot BUD13 protein sequences revealed notable associations. The sequences clustered into two main groups: (1) a major group containing all sequences derived from Poaceae species, including *P. notatum*; and (2) a minor group comprising sequences from the remaining species. When both LONG and SHORT protein variants were reported in the databases (*O. sativa*, *P. notatum*, *H. vulgare*, *Z. mays*, *E. curvula*, *T. turgidum*, *S. italica*, *A. officinalis*, and *Z. officinale*), they clustered together within the same branch, supporting the hypothesis that they originate from the same gene and represent splice variants or alternative transcription initiation sites (ATS). The *Paspalum* sequences clustered with those of related Poaceae species, such as *E. curvula*, *Panicum virgatum*, *S. italica and S. viridis*. *E. curvula* has been extensively studied in the context of apomixis, with numerous reports describing diplospory as the predominant reproductive mechanism and identifying candidate genes associated with the trait (Zappacosta *et al*., 2014, 2019). In contrast, research on apomixis in *P. virgatum* has been relatively limited, although moderate levels of apomictic reproduction have been proposed (McMillan and Weiler, 1959). At the genus level, one of the most extensively studied species, *P. maximum*, is known to exhibit both facultative aposporous apomixis and obligate sexual reproduction (Warmke, 1954; Savidan, 1982, 2019; Yamada-Akiyama *et al*., 2009; Bhandari *et al*., 2010; Kaushal *et al*., 2018). Regarding *S. italica*, studies on apomixis remain limited, although available evidence suggests a potential for apomictic reproduction (Emery, 1957; Fuyao *et al*., 2001). Furthermore, within the genus *Setaria*, apomixis has been documented in *S. viridis* (Dekker, 2004), and similar reproductive behavior has been proposed for *S. pumila*, *S. verticillata*, *S. leucopila*, *S. macrostachya*, and *S. texana*, all of which have been reported to produce both sexual and apomictic seeds (Dekker, 2004).

Remarkably, the *in situ* hybridization experiments reported here revealed differential expression patterns of *BUD13* transcript accumulation in sexual and apomictic ovules. At the premeiotic stage, signal was detected in the nucellus and integuments of ovules from both reproductive types, but was absent from the megaspore mother cell and the apospory initial cells (i.e., the cells that acquire a gametophytic fate) in aposporous ovules. This pattern was maintained during meiosis, as no detectable signal was observed in either the functional or the degenerating megaspores of the apomictic genotype, nor in the adjacent apospory initials. Furthermore, at anthesis, no *BUD13* signal was observed within the embryo sacs of aposporous plants, except in a few egg cells. During post-anthesis development, *BUD13* signal was clearly detected in parthenogenetically formed proembryos. Altogether, the results obtained in *P. notatum* suggest that *BUD13* is ubiquitously expressed throughout ovule development in sexual plants, whereas in apomictic plants it is downregulated in the female germline until the egg cell is formed. This downregulation may affect both the meiotic and aposporous germlines, which often coexist side by side during reproductive development. Once the mature embryo sac stage is reached, the egg cell of aposporous plants begins to express *BUD13*, and this expression increases during parthenogenetic embryo development, possibly because the gene is required for embryogenesis, as suggested by Xiong *et al*. (2019). Further investigation is needed to determine the basis of this differential *BUD13* expression in the female germline. In particular, it will be important to establish whether the presence of the extended 5′ UTR in the *SHORT* isoform from apomictic genotypes contributes to the selective suppression of this isoform in aposporous cell lineages.

## CONCLUSIONS

The central role of *BUD13* in splicing and its functional connection to *TGS1* make it a compelling candidate for the analysis of its role in apomixis development. Here, we provide evidence that *BUD13* is expressed as transcripts of different lengths in *P. notatum* and other monocots, possibly representing splice variants, which suggests a diversification of gene function. Moreover, we demonstrate structural and expression differences in *BUD13* activity between ovules of sexual and apomictic *P. notatum* plants, making *BUD13* a strong candidate for future functional analyses aimed at further elucidating its role in reproduction. We believe that this study contributes an additional piece to the intricate puzzle of developmental pathways governing maternal seed formation in plants.

## SUPPLEMENTARY INFORMATION

**Supplementary Data S1:** List of 110 BUD13 protein sequences retrieved from 40 monocot species and the two outgroup species *A. thaliana* and *S. cerevisiae*. Protein domains detected in the 110 BUD13 protein sequences and classification of each sequence as *LONG* or *SHORT*.

**Supplementary Fig. S1:** Pn_BUD13 protein alignment.

**Supplementary Fig. S2:** Phylogenetic analysis of 110 BUD13 protein sequences from monocots.

**Supplementary Fig. S3:** RNA in situ hybridization of *BUD13* transcripts in the reproductive organs of sexual (Q4188) and apomictic (Q4117) *P. notatum* individuals at the FM stage.

**Supplementary Fig. S4:** RNA in situ hybridization using the *BUD13* antisense probe (SP6) in ovaries of sexual (Q4188) and apomictic (Q4117) *P. notatum* at anthesis.

**Supplementary Fig. S5:** RNA in situ hybridization of *BUD13* transcripts in ovaries of sexual *P. notatum* (Q4188) at anthesis.

**Supplementary Fig. S6:** RNA *in situ* hybridization using the *BUD13* sense probe (T7) in adjacent sections of the same ovule from an apomictic *P. notatum* (Q4117) individual at anthesis.

## AUTHOR CONTRIBUTIONS

S.C.P., G.B., L.S. and J.P.A.O. contributed to the conception and design of the study. S.D., S.C.P., M.P., G.G. and F.P. performed sequence retrieval and bioinformatic analyses. L.S., S.D., M. S: V. and C.C. designed and cloned the probes and conducted the *in situ* hybridization experiments. L.S., S.D., J.P.A.O and M. S. V. carried out the qPCR validation. S.C.P. wrote the first draft of the manuscript. All authors reviewed and approved the final version of the manuscript.

## MATERIAL AND DATA AVAILABILITY STATEMENT

The plant material used in this study is available from Herbarium Carmen L. Cristobal, IBONE-CONICET-UNNE under the following voucher numbers: Q4117: CTES0541626; Q4188: CTES0542168. The datasets generated during and/or analyzed during the current study are available from the corresponding author on reasonable request.

## AI ASSISTANCE

English style, grammar and spelling were controlled with ChatGPT.

## ACKNOWLEDGEMENTS

This research was funded by the European Union’s Horizon 2020 Research and Innovation Program under the Marie Skłodowska-Curie Grant Agreements 872417 (MAD) and 101007438 (POLYPLOID). Additionally, funding was received by the National Agency for the Promotion of Research, Technological Development and Innovation, Argentina (PICT 2021-539) and National University of Rosario, Argentina (PID 800 20220700266UR). LS, MP, JPAO and SCP are research staff members of CONICET, Argentina.

