## Supplementary figures for "The BUD13 splicing regulator: transcript structure and expression in ovules of sexual and apomictic *Paspalum notatum*"

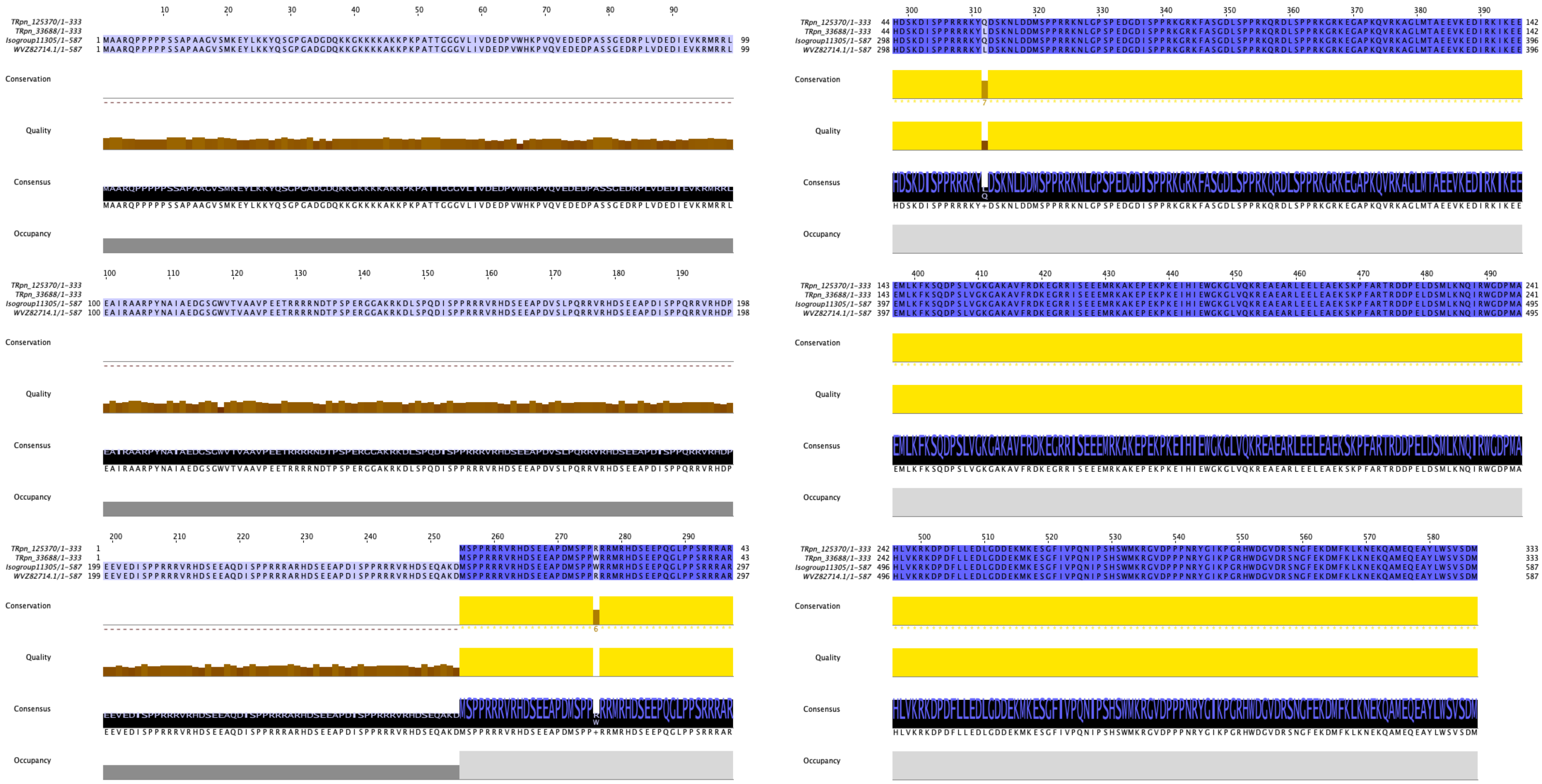

**Supplementary Fig. S1: Pn_BUD13 protein alignment.** Alignment of the full-length BUD13 proteins predicted from 1) the genomic locus PnR1_U9M48_029946 (WVZ82714.1) (Vega et al., 2024); 2) the Roche 454 floral transcriptome (isogroup 11305, LONG) (Ortiz et al., 2017); and the Illumina HiSeq floral transcriptome TRpn_125370 (SHORT1) and TRpn_33688 (SHORT2) (Podio et al., 2021).

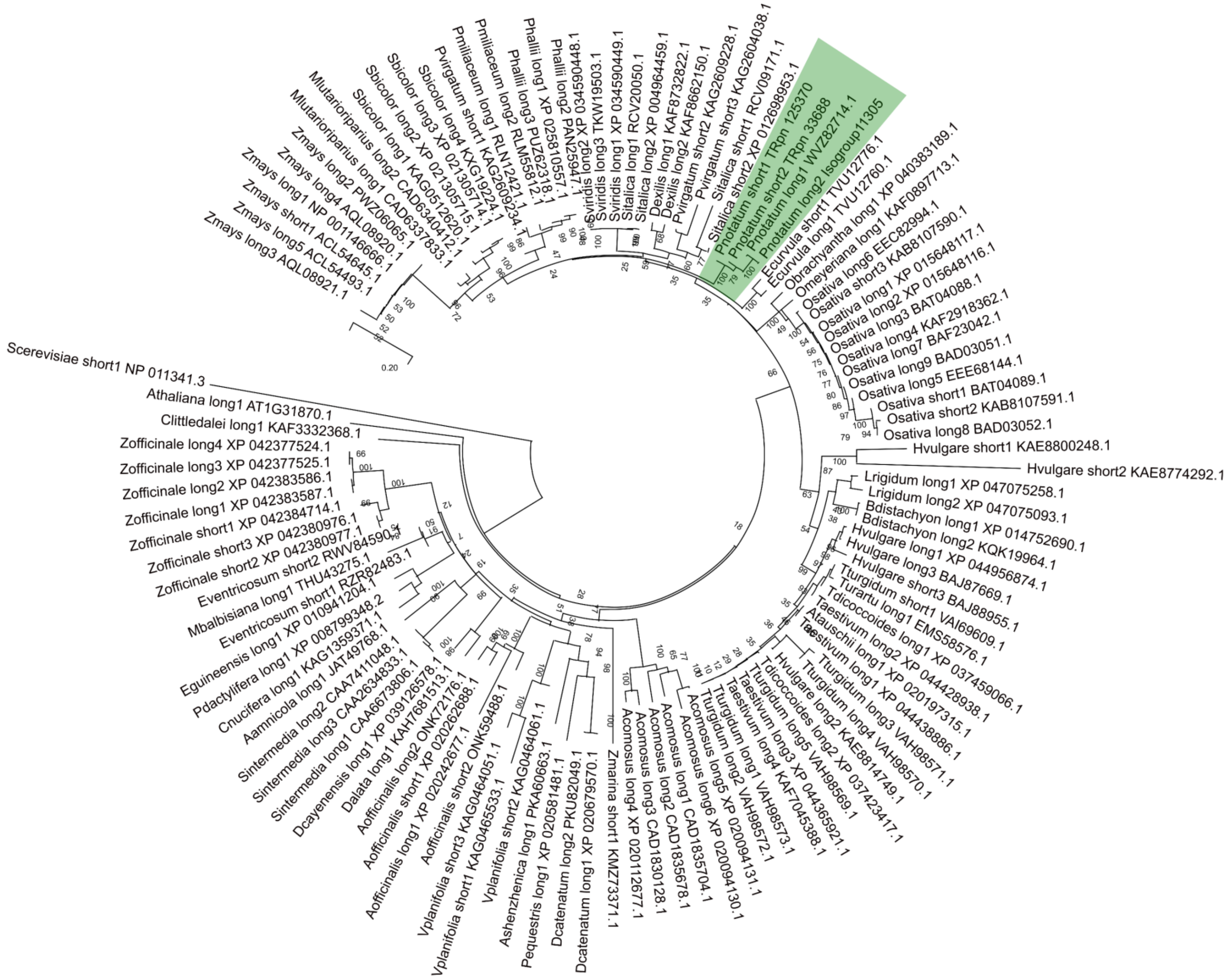

**Supplementary Fig. S2: Phylogenetic analysis of 110 BUD13 protein sequences from monocots.** The *Paspalum notatum* sequences (marked in green) grouped within the Poaceae cluster.

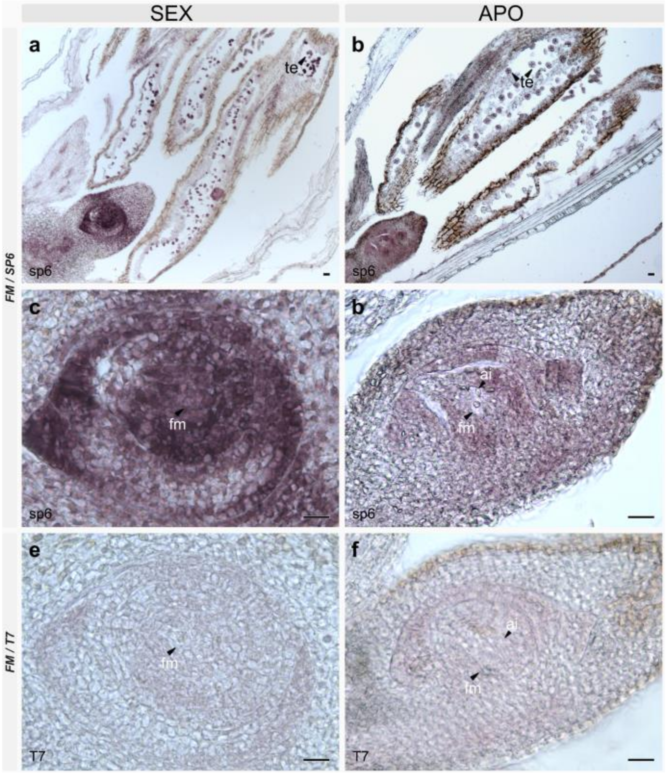


**Supplementary Fig. S3. RNA in situ hybridization of BUD13 transcripts in the reproductive organs of sexual (Q4188) and apomictic (Q4117) *P. notatum* individuals at the FM stage.** (a–d) Hybridization with the antisense probe (SP6) in sagittal sections of ovules from sexual (a, c) and apomictic (b, d) plants. Panels (a, c) and (b, d) show different focal planes of the same section. (e, f) Hybridization with the sense probe (T7) in sagittal sections of ovules from sexual (e) and apomictic (f) individuals. ai: apospory initials; apo: ovules of Q4117; fm: functional megaspore cell; te: tetrads; sex: ovules of Q4188. Bars: 10 µm.

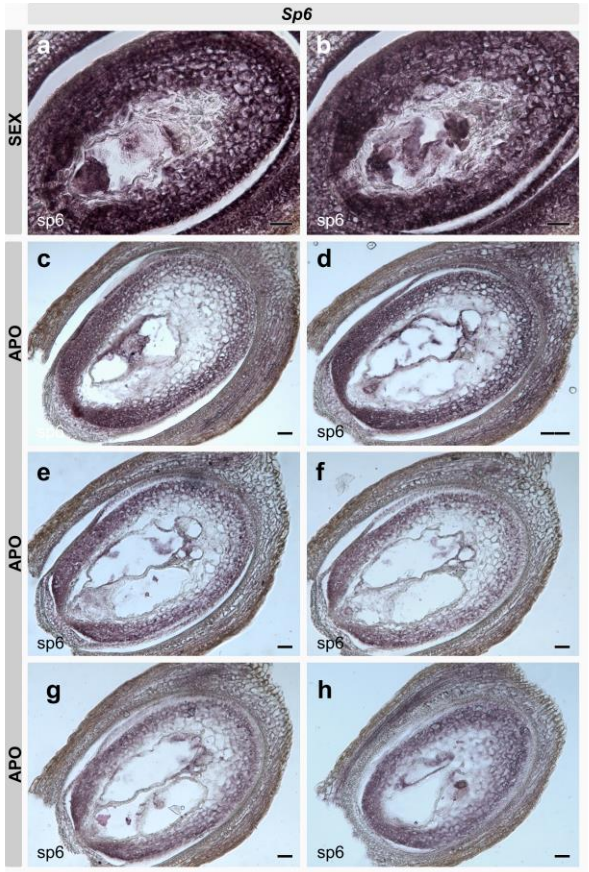


**Supplementary Fig. S4. RNA in situ hybridization using the BUD13 antisense probe (SP6) in ovaries of sexual (Q4188) and apomictic (Q4117) *P. notatum* at anthesis.** (a, b) Adjacent sections of the same ovule shown in Fig.3, from a sexual individual. (c–h) Adjacent sections of the same ovule shown in Fig.3, from an apomictic individual. Bars: 10 µm.

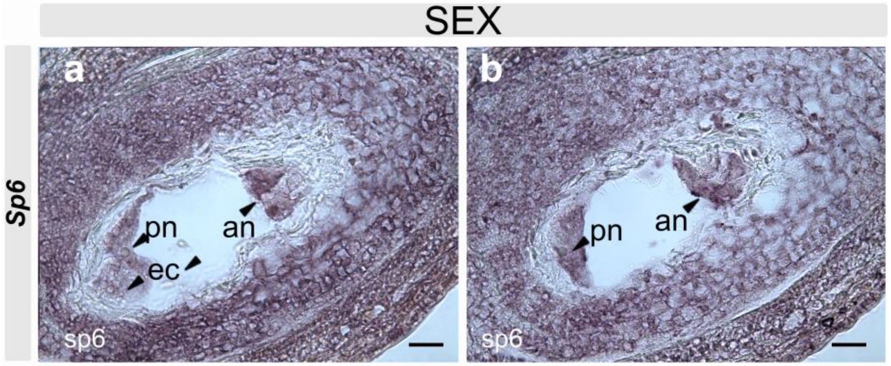


**Supplementary Fig. S5. RNA in situ hybridization of BUD13 transcripts in ovaries of sexual *P. notatum* (Q4188) at anthesis.** (a, b) Hybridization with the BUD13 antisense probe (SP6) in adjacent sections of the same ovule. an: antipodal cells; ec: egg cell; pn: polar nuclei. Bars: 10 µm.

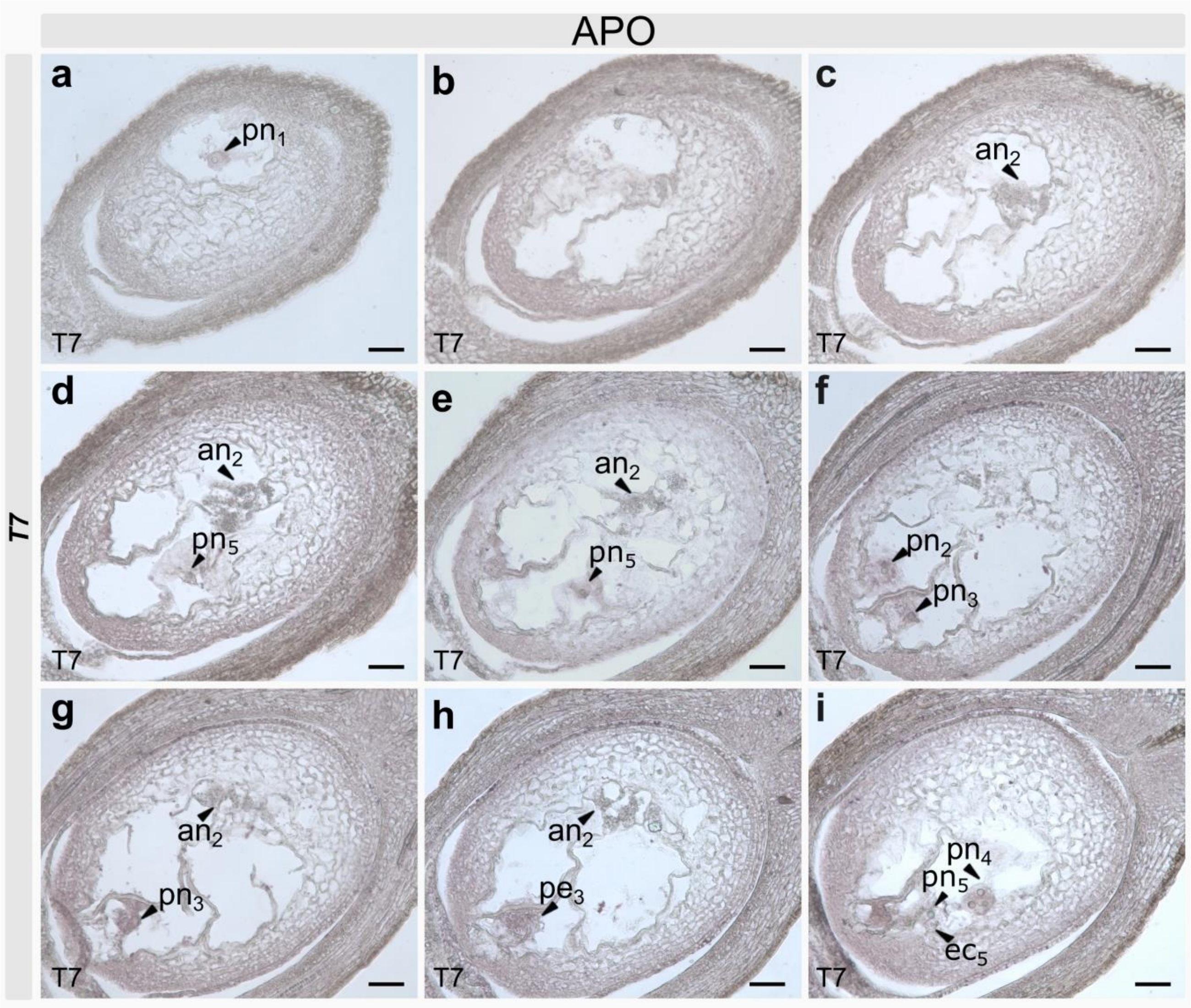


**Supplementary Fig. S6. RNA in situ hybridization using the BUD13 sense probe (T7) in adjacent sections of the same ovule from an *apomictic P. notatum* (Q4117) individual at anthesis.** an: antipodal cells; es: embryo sac; pe: pro-embryo. Subscript numbers indicate the embryo sac in which each structure is located. Bars: 10 µm.
